# Deltamethrin contact exposure mediated toxicity and histopathological aberrations in tissue systems of cockroach species *Periplaneta americana* and *Blatella germanica*

**DOI:** 10.1101/2021.12.06.471460

**Authors:** Sunil Dhiman, Kavita Yadav, BN Acharya, DP Nagar, Rama Rao Ghorpade

## Abstract

Cockroach species *Periplaneta americana* and *Blatella germanica* potentially survive in locations close to human activity. Besides spoiling food material, cockroaches also transfer pathogens of different diseases among human. Since the insecticides have been used extensively to control cockroaches, information on their insecticide susceptibility and toxicity at cellular level may be crucial. In the study, deltamethrin toxicity as well as the deltamethrin-mediated cytomorphological changes in brain, ovary and midgut of the two important cockroach species has been assessed. Different concentrations [0.00025% (0.0025mg/ml), 0.0025% (0.025mg/ml), 0.025 (0.25mg/ml), 0.25% (2.5mg/ml), 0.5% (5mg/ml), 1% (10mg/ml)] of deltamethrin in acetone were used to expose test species in WHO bottle assay. Knockdown was recorded after 5 min interval while delayed mortality was observed after 24 hr. Brain, ovary and gut were dissected post 1 hr exposure and 24 hr holding (for 0.25%, 0.5% and 1% concentration), and tissues were processed for microscopic analysis. Deltamethrin exposed cockroaches and dissected tissues were used to estimate deltamethrin using HPLC. At 0.00025% (lowest concentration), the percentage knock-down observed was 66.7% for *P. americana* and 80% *B. germanica* respectively (R^2^= 0.78; p=0.0001) in 1 hr. KDT_50_ value was found to be 8.7 min (95% CI: 7.3-10.2), while KDT_99_ was 20.7 min (95% CI: 16.0-35.7) in *P. americana* at 1% concentration. Whereas, the KDT_50_ and KDT_99_ values for *B. germanica* were 7.4 min (95% CI: 5.4-9.1) and 27.4 min (95% CI: 18.2-80.0) at similar concentration. LD_50_ and LD_95_ values (for 60 min standard exposure) were 0.0006 % (95% CI: 0.00-0.001) and 0.034% (95% CI: 0.013-0.49) respectively for *P. americana*, while these values were 0.0005 (95% CI: 0.00-0.001) and 0.04 (95% CI: 0.01-0.23) for *B. germanica*. Exposure to 1% deltamethrin induced considerable toxic effect in the epithelial cells in the midgut. HPLC estimated 0.21±0.05 mg (95% CI - 0.18-0.25; CoV 23.9%) deltamethrin in *P. americana* post 1% exposure. Even short term exposure of low concentration of synthetic pyrethroid deltamethrin displayed immediate knockdown and delayed mortality in both the test species. Considerable histological damage was observed in both the insects at 1% exposure. In India, resistance to deltamethrin may have been reported among different insects due its extensive use, however the formulations such as insecticide paints, attractant baits etc. developed using deltamethrin as active ingredient could be useful in cockroach control operations.

## Introduction

Both *Periplaneta americana* and *Blatella germanica* species of cockroaches are important peridomestic pests in urban communities in majority of Asian countries. These potentially survive in locations close to humans or human activity. Therefore this arthropod pest is primarily found in residential areas, sewage system, farm produce markets, grain stores, trains and different commercial establishments. Cockroaches are gregarious and it has been found that different species are able to aggregate at the same location [1].

*P. americana* (American cockroach) is generally reddish brown, large in size and may measure upto 34-53 mm in length. This species is highly mobile and studies conducted have indicated the movement to several hundred meters through sewer systems and channels into the neighboring homes. On the other hand *B. germanica* (German cockroach) is a small brown to black coloured species and may measure about 11-16 mm in length [2]Weaving et al. 2003) long. Of the few cockroach species that are domestic pests, it probably is the most widely troublesome of all [3]Bonnefoy et al. 2008).

Cockroaches have been regarded as major pest species which are both a nuisance and can cause severe health problems. These not only spoil the food materials, but also transfer pathogens of different diseases and may cause allergic reactions, and in some cases psychological distress too [4]Brenner 1995). Although cockroach’s role in direct transmission of pathogen has not been much established, but the studies have shown that many pathogenic organisms such as poliomyelitis viruses, protozoa, bacteria, fungi, and helminthes have been associated with the cockroaches [5-7]Fathpour et al. 2003; Saichua et al. 2008; Tatfeng et al. 2005). These have been reported as vectors of nosocomial infections, while both these species have been found carrying antibiotic resistance bacteria [8,9]Pai et al. 2004, 2005). *B. germanica* regularly inhabits the food preparation areas during night thus such areas may become contaminated. Study by Tachbele et al. [10]2006) has identified a species of *Salmonella, Shigella flexneri, Escherichia coli, Staphylococcus aureus* and *Bacillus cereus* from *B. germanica* in Ethopia. These studies indicate that cockroach species may act as possible reservoir and vector of many pathogens of human importance. In addition to this, both these species of cockroaches cause allergies to human [11]William, 2016). The major allergens, Bla g1, Bla g2 and Per a1, have been found in the saliva, fecal material, secretions, cast skins and debris. Studies have also demonstrated the relationship between cockroach exposure and poor asthma outcomes among those who are exposed to high levels of cockroach allergens [12](Do et al. 2016). Therefore the households can have allergic reactions due to exposure to cockroach body parts and feces, which in most of the cases are asthmatic, but can be life threatening in some cases.

The effective control of cockroaches can reduce the diseases associated, allergen spread and associated mortality and morbidity among humans [13](Rabito et al. 2017). The control of cockroaches mostly relies on using insecticides, however it is crucial to delineate the effectiveness of these insecticides against different cockroach species in an area of interest. At present, the extensive use of different insecticide groups in the control of virtually all harmful arthropod pests has raised serious concerns about their efficacy at a relatively low and sustainable concentration. Like many arthropod vectors, cockroaches have shown resistance to different insecticides [14,16,15](Gondhalekar et al. 2012; Pai et al. 2005; Zhu et al. 2016). This may be because cockroaches live in relatively close and large populations, thus facilitate rapid selection for different insecticides they are exposed. However another study [17](Syed et al. 2014) has shown that many synthetic pyrethroids were effective against American cockroaches. The study further suggested that insecticides efficacy differ among the locations. Therefore estimation of insecticide susceptibility of cockroach species to prominently used insecticides may guide substantial use of such insecticide for the control in an area of interest.

The present study was undertaken with an objective to understand the effectiveness of synthetic pyrethroid deltamethrin against two prominent cockroach species *P. americana* and *B. germanica* under laboratory conditions. The study presents an effort to generate deltamethrin sensitivity data and to assess whether deltamethrin as single active ingredient could be used in effective control of cockroaches.

## Methods

### Test organism

The culture of both the cockroach species are maintained in the insectary of Defence Research and Development Establishment, Gwalior. However for the present study the cockroach species were collected from the kitchens, basements, cupboards and related areas in Gwalior, Madhya Pradesh (India) (Latitude: 26° 13’ 5.8332’’ N and longitude: 78° 10’ 58.1916’’ E) and the colonies were maintained in 3 Lt capacity glass jar fitted with specially designed metal steps inside the jar for movement (Figure 1) at 25±2 °C temperature, 60±5% relative humidity and exposed to a photoperiod of 12:12 (L:D) [17](Snoody et al. 2014). The insects were provided water, wheat flour and powdered dog biscuits *ad libitum*. Adult male cockroaches [19,18](WHO, 1970; Snoody et al. 2014) of 5-10 days old (almost equal size) from the reared generations were used for the experiments.

**Figure 1:**
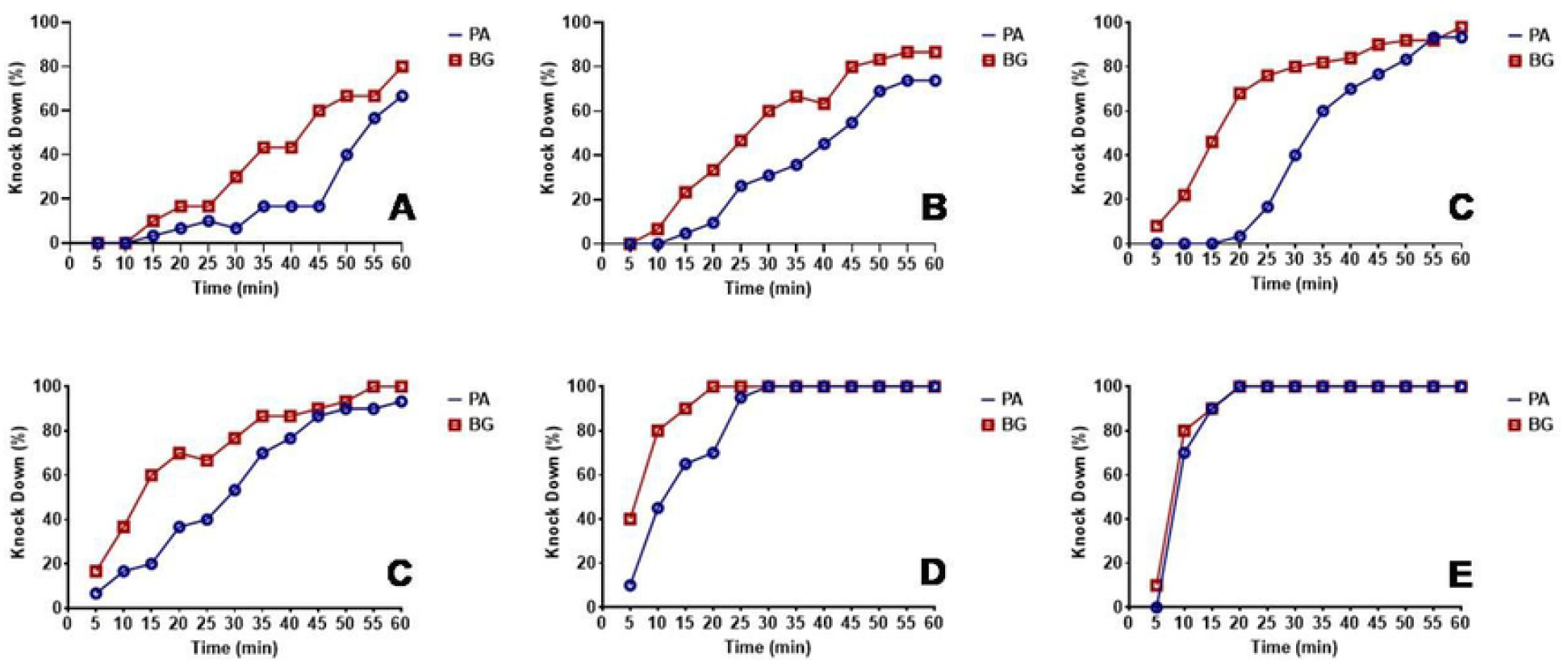
Knock-down rate for *P. americana* (PA) and *B. germanica* (BG) for different concentrations of deltamethrin. (A) 0.00025%, (B) 0.0025%, (C) 0.025%, (D) 0.25%, (E) 0.5%, (F) 1%.

### Knockdown and mortality bioassays

Technical grade deltamethrin (98% purity) obtained from M/S Tagros Chemical India, Chennai was used for the experimental exposure. The insecticide in different concentrations [0.00025% (0.0025mg/ml), 0.0025% (0.025mg/ml), 0.025 (0.25mg/ml), 0.25% (2.5mg/ml), 0.5% (5mg/ml), 1% (10mg/ml)] were made in acetone. Two ml of concentration was applied uniformly on the surface of glass bottle jar (500 ml capacity; area 487.8 cm^2^) and left to dry for 1 hr in a fume hood. For each experiment five adult cockroaches were exposed for 1 hr, and at least 30 cockroaches were taken for each exposure. In the control treatment, only acetone was used. Knockdown was observed after 5 min interval. After the exposure, the cockroaches were transferred to holding plates and provided with ample of food and water. Mortality was recorded after 24 hr, 48 hr and 72 hr post exposure. Exposure lethal concentrations (LC) were determined by exposing the cockroaches at different concentrations for a given time; whereas lethal time (LT) was estimated by exposing the cockroaches at a given concentration for different time periods. Petroleum jelly mixed with mineral oil was thin layer coated at the open end of the jar to prevent the escape of insects during experiments. The experiments were performed as described previously [20,18,19](Scharf et al. 1995; Snoody et al 2014; WHO, 1970).

### Histological analysis

For histopathological evaluation, cockroach species *P. americana* exposed to 0.25% (group I; N=5), 0.5% (group II; N=5), and 1% (group III; N=5) of deltamethrin for 1 hr. Whereas the insects exposed to acetone (group IV; N=5) were taken as control. Brain, ovary and gut were dissected in saline solution (0.13 M NaCl; 0.01 M; Na_2_HPO_4_ 0.02 M; KH_2_PO_4_; pH 7.2) post 24 hr of exposure and fixed in 10% neutral buffered formalin solution for another 24 hr at room temperature. After fixation and overnight washing in distilled water, the extracted tissues were processed for dehydration in graded series of alcohol and toluene in auto-tissue processor (Leica TP-1020, Germany). Processed tissues were embedded in paraffin wax (Leica, Germany). Multiple sections of 2.5-3 µm thickness from each block were cut on rotatory microtome (Thermo Scientific, Microm, USA), mounted on pre-coated glass slide and air dried overnight. The sections were deparaffinised and stained with haematoxylin and eosin [21](McMannus and Mowry, 1965) in auto-stainer (Leica, Germany) and cover slipped by auto cover slipper (Leica, Germany). After drying, sections were photographed under light microscope (Leica DMLB, Germany) using DM-500 camera (Leica, Germany).

### HPLC estimation of deltamethrin

*P. americana* adult exposed to 1% deltamethrin were weighted and taken (N=10) for HPLC quantification of deltamethrin on cockroach surface. Whereas, the brain, gut and ovary tissues dissected from the cockroaches exposed to 0.25%, 0.5% and 1% deltamethrin were used for estimation. For quantification from whole cockroach, each sample was taken into 5 ml ice-cold acetonitrile (HPLC grade), homogenized for 10 min and sonicated for 15 min. Similarly internal tissues were homogenized in 500 µl acetonitrile, sonicated for 15 min and filtered before using for HPLC. The samples were filtered through 0.22 µ PVDF membrane filter and stored at -4°C for use in HPLC. Active compound deltamethrin was separated on a C18 reverse-phase column (4.6×250 mm, particle size 5 μm; Waters XTerra™) maintained at room temperature (25°C). Methanol and water (80:20 v/v) were used as mobile phase with flow rate of 1 mL/min using binary pump (Waters, 1225). Detection was performed using dual absorbance detector (Waters, 2487) at 280 nm. Prepared samples were injected through Rheodyne injector (injection volume 20μL). The HPLC instrument used was controlled by Empower computer programme (Europa Science, Ltd., Cambridge, UK) for data collection and overall instrument control during the experiments.

### Statistical analysis

Knockdown at different time intervals was observed and presented in percent knockdown (%KD), whereas mortality was scored after 24 hr of exposure and presented as corrected mortality. Spearman correlation was performed to assess the knockdown for both the species at different interval, while using that the data did not follow Gaussians distribution. Log dose probit method has been used to estimate the KD, LD and LT values using LdP, Software (Ihabsoft, Turkey). Results were considered significant when p<0.05 at 95% confidence intervals.

## Results

### Knock-down bioassays

The percent knock-down achieved at different time intervals in cockroach species *P. americana* and *B. germanica* after 1 hr exposure to different concentrations of deltamethrin has been depicted in fig. 1, whereas the overall knock-down effect observed has been presented in Table 1. At lowest concentration of 0.00025%, the percentage knock-down observed was 66.7% for *P. americana* and 80% *B. germanica* respectively (R^2^= 0.78; p=0.0001) in 1 hr. On the other hand after 20 min during the exposure at 1% deltamethrin concentration, both the species exhibited 100% knockdown (Fig. 1). The KDT_50_ value was 8.7 min (95% CI: 7.3-10.2) whereas KDT_99_ was 20.7 min (95% CI: 16.0-35.7) in *P. americana* at 1% concentration. Nevertheless both KDT_50_ and KDT_99_ values were found increased at 0.00025% and recorded to be 57.7 min (95% CI: 51.4-69.4) and 219.2 min (95% CI: 147.2-459.7) respectively. On the other hand for *B. germanica* the KDT_50_ and KDT_99_ values were 7.4 min (95% CI: 5.4-9.1) and 27.4 min (95% CI: 18.2-80.0) at 1% concentration, while found to be 39.8 min (95% CI: 36.0-44.4) and 176.4 min (95% CI: 126.3-308.5) at 0.00025% respectively (Table 1). The probit analysis used for determining the knock-down percent at different time intervals for all the treatments displayed normal distribution and did not deviate from the linearity (p≤0.11; □^2^≥13.2).

**Table 1:** Deltamethrin exposure knock-down values for P. americana and B. germanica.

| Species | Concentration (%) | KDT<sub>50</sub><br>(95% CI) | KDT<sub>99</sub><br>(95% CI) | p (Z) | Slope (+/-) |
| --- | --- | --- | --- | --- | --- |
| *P. americana* | 0.00025 | 57.7<br>(51.4-69.4) | 219.2<br>(147.2-459.7) | 0.11 (13.2) | 4.0 (+/-0.6) |
|  | 0.0025 | 40.4<br>(37.4-43.8) | 156.2<br>(120.9-229.8) | 0.96 (2.5) | 3.9 (+/-0.4) |
|  | 0.025 | 39.0<br>(35.6-42.7) | 138.6<br>(104.5-225.4) | 0.51 (6.3) | 4.2 (+/-0.6) |
|  | 0.25 | 23.7<br>(20.8-26.5) | 131.5<br>(100.2-195.8) | 0.37 (10.9) | 3.1 (+/-0.3) |
|  | 0.5 | 11.5<br>(9.2-13.9) | 53.3<br>(35.6-122.8) | 0.52 (2.3) | 3.5 (+/-0.7) |
|  | 1 | 8.7<br>(7.3-10.2) | 20.7<br>(16.0-35.7) | 0.44 (0.6) | 6.2 (+/-1.3) |
| *B. germanica* | 0.00025 | 39.8<br>(36.0-44.4) | 176.4<br>(126.3-308.5) | 0.93 (3.1) | 3.6 (+/-0.5) |
|  | 0.0025 | 26.5<br>(23.5-29.5) | 135.0<br>(102.1-206.5) | 0.99 (1.8) | 3.3 (+/-0.4) |
|  | 0.025 | 16.0<br>(14.0-18.0) | 116.7<br>(92.0-160.5) | 0.68 (7.5) | 2.7 (+/-0.2) |
|  | 0.25 | 13.7<br>(10.5-15.7) | 128.6<br>(87.8-235.5) | 0.98 (2.0) | 2.4 (+/-0.3) |
|  | 0.5 | 8.9<br>(5.9-9.6) | 29.6<br>(17.8-167.0) | 0.76 (0.09) | 3.3 (+/-0.9) |
|  | 1 | 7.4<br>(5.4-9.1) | 27.4<br>(18.2-80.0) | 0.93 (0.00) | 4.1 (+/-0.9) |

KDT = knockdown time in minutes; CI = confidence interval; p significant if <0.05.

### Time-concentration mortality

The exposure time mortality data has been shown in table 2. It was found that post 10 min exposure of deltamethrin and 24 hr holding the LD_50_ and LD_95_ values were 1.06 % (95% CI: 0.44-7.07) and 117.15% (95% CI: 13.18-62099.0) respectively, while for 60 min standard exposure these values were found to be 0.0006 % (95% CI: 0.00-0.001) and 0.034% (95% CI: 0.013-0.49) respectively for *P. americana* species. Furthermore LD_50_ and LD_95_ values for *B. germanica* species were found to be 0.35% (0.16-1.60) and 100.6 (9.2-309939.6) after 10 min exposure while 0.0005 (95% CI: 0.00-0.001) and 0.04 (95% CI: 0.01-0.23) after 60 min exposure respectively. Furthermore the mortality values obtained 48 h and 72 holding post 1 hr exposure have been displayed in table 3. It was observed that the LD_99_ values were 0.01% (p=0.88) and 0.02% (p=0.27) respectively for *P. americana* and *B. germanica* species post 72 of exposure (table 3). Probit model used to determine the lethal time values by using exposure for different time periods at a concentration showed that the LT_50_ and LT_99_ values were 72.5 min and 199.6 min respectively for *P. americana* at lowest concentration used (Table 4). Similarly, these values were 75.9 min and 380.8 min respectively for *B. germanica* at similar concentration. It was found that the corrected mortality after 24 hr, 48 hr and 72 hr holding time post 1 hr exposure was 100% each in 0.25% deltamethrin in both the species, whereas varied from 40% to 70% in *P. americana* and 43% to 70% in *B. germanica* for 0.00025% deltamethrin (Supplementary file 1).

**Table 2:** Lethal dose (LD) values of P. americana and B. germanica for deltamethrin exposure to different time periods

|  | Exposure time (min) | LD50 (95% CI) | LD95 (95% CI) | Slope (+/-) | 檢(p) | R |
| --- | --- | --- | --- | --- | --- | --- |
| *P. americana* | 10 | 1.06<br>(0.44-7.07) | 117.15<br>(13.18-62099.0) | 0.80 (0.20) | 0.80 (0.2) | 0.94 |
|  | 20 | 0.29<br>(0.15-0.61) | 11.29<br>(3.27-151.82) | 1.03 (0.20) | 2.50 (0.5) | 0.97 |
|  | 30 | 0.07<br>(ND) | 3.8<br>(ND) | 0.95 (0.22) | 6.81 (0.03) | 0.92 |
|  | 40 | 0.06<br>(0.03-0.17) | 3.44<br>(0.72-97.88) | 0.92 (0.18) | 1.93 (0.4) | 0.98 |
|  | 50 | 0.002<br>(0.00-0.005) | 0.38<br>(0.088-7.67) | 0.71 (0.14) | 0.17 (0.9) | 1 |
|  | 60 | 0.0006<br>(0.00-0.001) | 0.034<br>(0.013-0.49) | 0.92 (0.20) | 1.79 (0.2) | 0.97 |
| *B. germanica* | 10 | 0.35<br>(0.16-1.60) | 100.6<br>(9.2-309939.6) | 0.67 (0.19) | 3.10 (0.2) | 0.89 |
|  | 20 | 0.07<br>(0.04-0.13) | 4.18<br>(1.49-23.36) | 0.92 (0.13) | 7.76 (0.1) | 0.96 |
|  | 30 | 0.02<br>(0.01-0.03) | 2.97<br>(0.79-31.3) | 0.72 (0.11) | 3.94 (0.3) | 0.97 |
|  | 40 | 0.01<br>(0.003-0.01) | 2.1<br>(0.53-24.43) | 0.66 (0.10) | 1.51 (0.7) | 0.98 |
|  | 50 | 0.002<br>(0.00-0.005) | 0.41<br>(0.13-2.68) | 0.72 (0.11) | 3.97 (0.3) | 0.99 |
|  | 60 | 0.0005<br>(0.00-0.001) | 0.04<br>(0.01-0.23) | 0.89 (0.17) | 1.91 (0.2) | 0.97 |

**Table 3:** Mortality effect on P. americana and B. germanica species post 24 hr, 48 hr and 72 hr after 1 hr deltamethrin exposure

| Species | Time | L D<sub>50</sub> (95% CI) | L D<sub>99</sub> (95% CI) | p ( $\chi^2$ ) |
| --- | --- | --- | --- | --- |
| PA | 24 h | 0.0006 (0.0002-0.0013) | 0.21 (0.44-8.93) | 0.2 (1.79) |
|  | 48 h | 0.0002 (0.000-0.0004) | 0.04 (0.01-1.94) | 0.3 (0.99) |
|  | 72 h | 0.00 (ND) | 0.01 (ND) | 0.9 (0.02) |
| BG | 24 h | 0.0005 (0.0002-0.0011) | 0.21 (0.54-3.48) | 0.2 (1.91) |
|  | 48 h | 0.0003 (0.0001-0.0006) | 0.04 (0.01-0.52) | 0.7 (0.12) |
|  | 72 h | 0.00 (ND) | 0.02 (ND) | 0.3 (1.21) |

L D<sub>50/99</sub> - Lethal dose 50%/99%; hr - hours; CI - confidence interval; p significant if < 0.05; ND - Not determined as value of g > 0.4. PA- *P. americana*, BG - *B. germanica*.

**Table 4:** Lethal time (LT) values of P. americana and B. germanica for different concentration of deltamethrin exposure

|  | Exposure concentration (%) | LT<sub>50</sub><br>(95% CI) | LT<sub>99</sub><br>(95% CI) | Slope (+/-) | Chi (p) | R |
| --- | --- | --- | --- | --- | --- | --- |
| *P. americana* | 0.00025 | 72.5<br>(65.0-94.9) | 199.6<br>(131.0-658.5) | 5.3 (1.3) | 2.2 (0.3) | 0.9 |
|  | 0.0025 | 53.6 (ND) | 236.6 (ND) | 3.6 (0.8) | 17.3 (0.001) | 0.81 |
|  | 0.025 | 44.1 (ND) | 205.1 (ND) | 3.5 (0.6) | 24.7 (0.0001) | 0.8 |
|  | 0.25 | 21.9<br>(16.0-27.7) | 168.6<br>(92.6-733.5) | 2.6 (0.6) | 4.0 (0.3) | 0.9 |
|  | 0.5 | 13.3<br>(9.6-16.3) | 47.7<br>(34.4-130.0) | 4.2 (1.0) | 3.2 (0.07) | 0.91 |
| *B. germanica* | 0.00025 | 75.9<br>(59.6-152.6) | 380.8<br>(175.9-4891.6) | 3.3 (0.9) | 1.5 (0.7) | 0.97 |
|  | 0.0025 | 46.6<br>(40.2-57.8) | 231.0<br>(135.6-824.3) | 3.4 (0.7) | 0.1 (0.99) | 1 |
|  | 0.025 | 25.7 (ND) | 181.4 (ND) | 2.7 (0.3) | 18.7 (0.0009) | 0.89 |
|  | 0.25 | 15.1<br>(10.7-18.8) | 119.0<br>(74.6-310.0) | 2.6 (0.5) | 0.6 (0.9) | 0.99 |
|  | 0.5 | 7.0 (ND) | 54.7 (ND) | 2.6 (1.0) | 0.05 (0.8) | 1 |

LT<sub>50/99</sub> - Lethal time 50/99%; CI - confidence interval; p significant when <0.05. ND - Not determined as value of $g > 0.4$ .

### Histological analysis

#### Brain

The brain of cockroach species *P. americana* consisted of well defined regions having condensed neuropiles in the central region while neural cell bodies in periphery. The histological examination of brain tissue from control and treatment did not show any considerable differences and were normal in histology with intact neurons. There was no abnormality seen in the treatment samples compared to the control (Fig 2. A-D).

**Figure 2:**
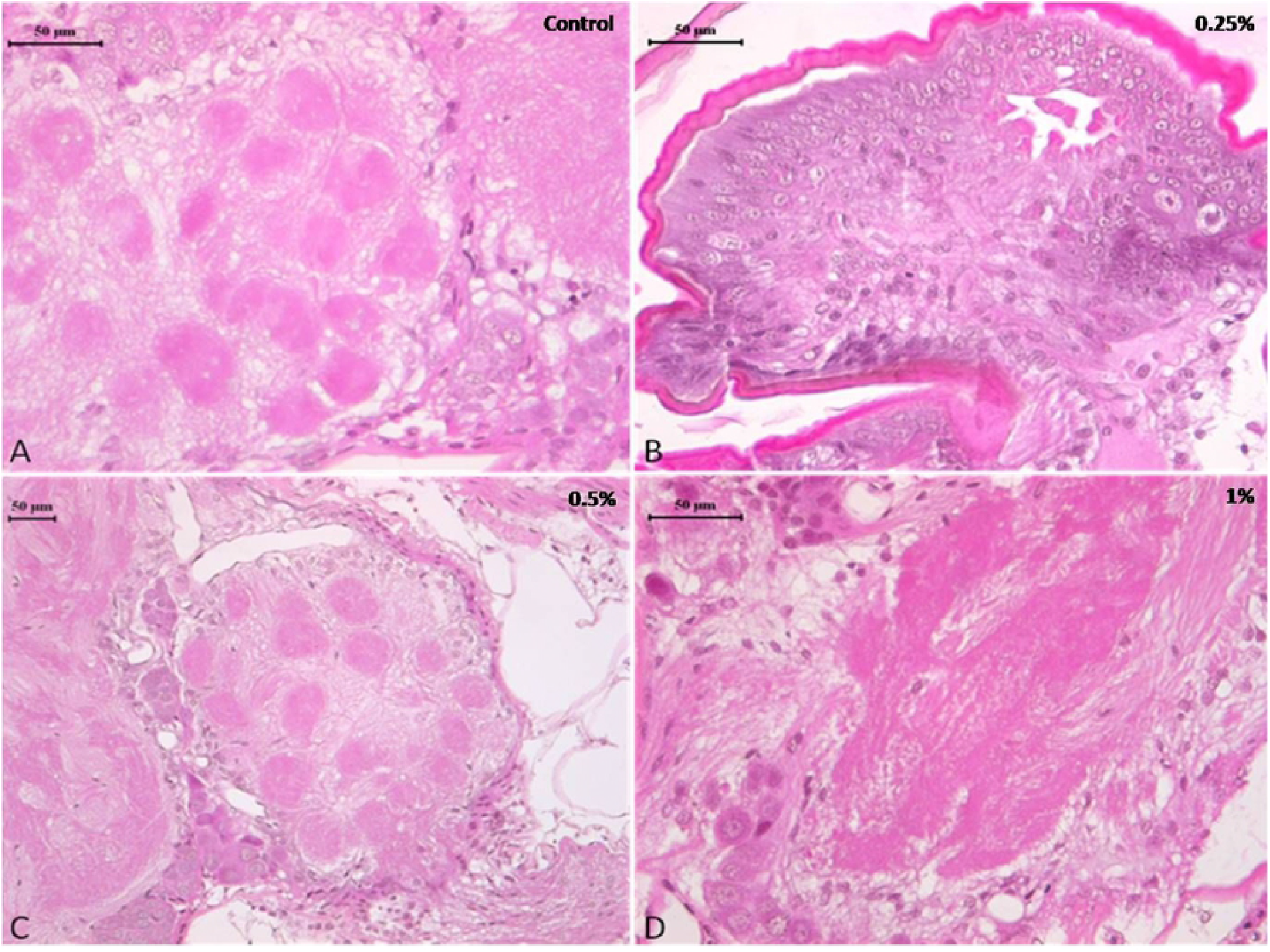
Photomicrograph of control and deltamethrin treated cockroach brain: A, Control; B, treated with 0.25% deltamethrin; C, treated with 0.5% deltamethrin; D, treated with 1% deltamethrin.

#### Ovary

The histopathological examination of ovary of both control as well as deltamethrin exposed (0.25%, 0.5% and 1%) cockroach species *P. americana* showed normal ovarian follicles with no deteriorating effects. The developing ovarian follicles exhibited normal development and yolk construction. Yolk material in oocytes was homogenous and the yolk bodies were surrounded by clear rings, with no cracks and fissures (Fig 3 A-D).

**Figure 3:**
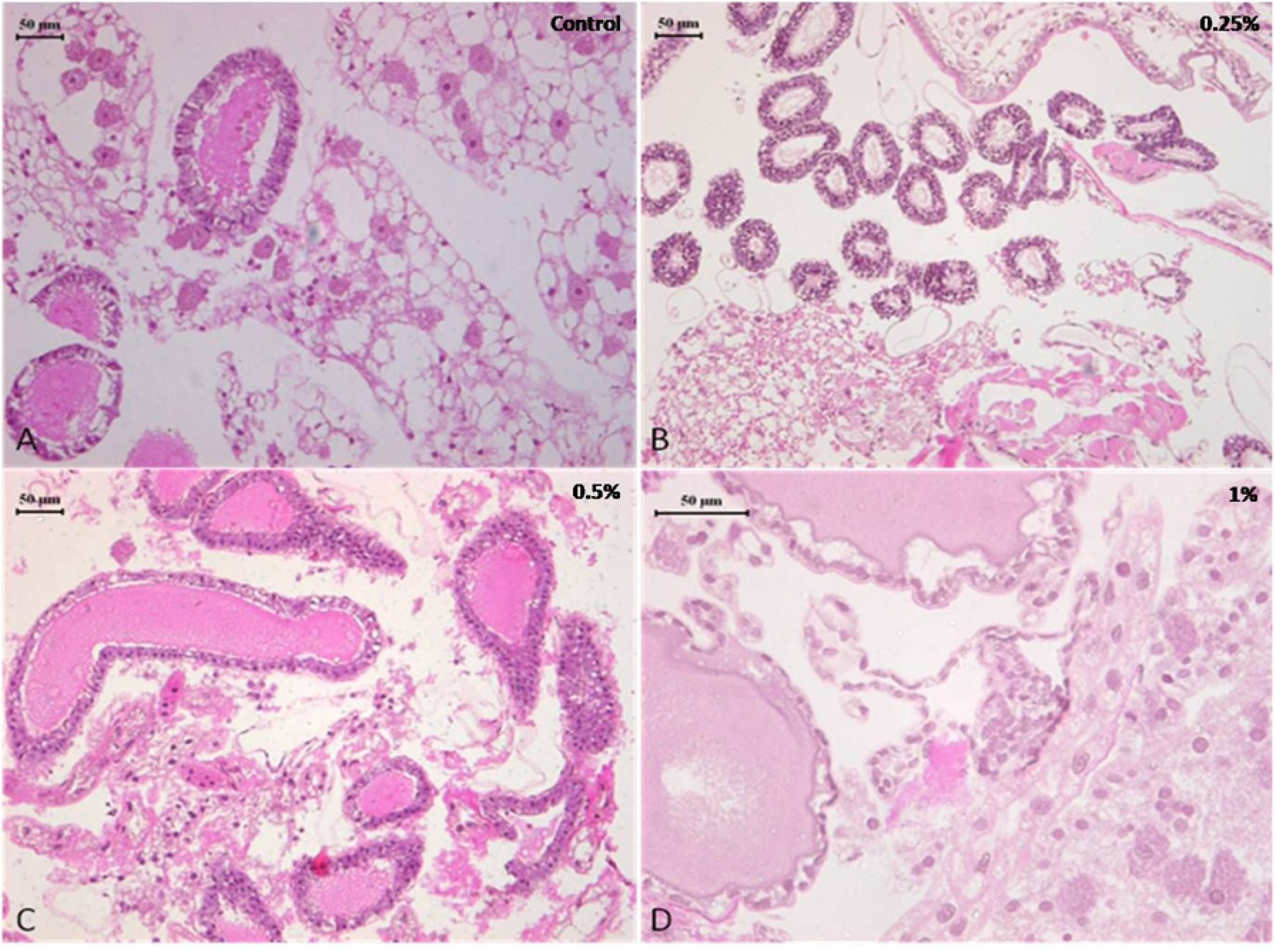
Photomicrograph of control and deltamethrin treated cockroach ovary: A, Control; B, treated with 0.25% deltamethrin; C, treated with 0.5% deltamethrin; D, treated with 1% deltamethrin.

#### Midgut

The midgut of control and 0.25% and 0.5% deltamethrin treated cockroaches displayed normal epithelium cells having a striated border with well defined nucleus. The epithelium cells rest upon a basement membrane followed by a normal inner layer circular muscles and a regular outer layer of longitudinal muscles. Midgut contained normal peritrophic membrane within its central lumen and is present as a thin transparent membrane. The histology of midgut of group I and II insects were normal and did not show considerable difference to the control (Fig 4 A-C). After treatment with 1% of deltamethrin (group III) disorganization and disintegration of epithelial cells was observed in the midgut of cockroach (Fig. 4D). Degenerated cytoplasm with distortion in circular and longitudinal muscle, disappearance of cell boundaries of the epithelial cells and vacuolization were some common histopathological changed seen in group III as compared to control.

**Figure 4:**
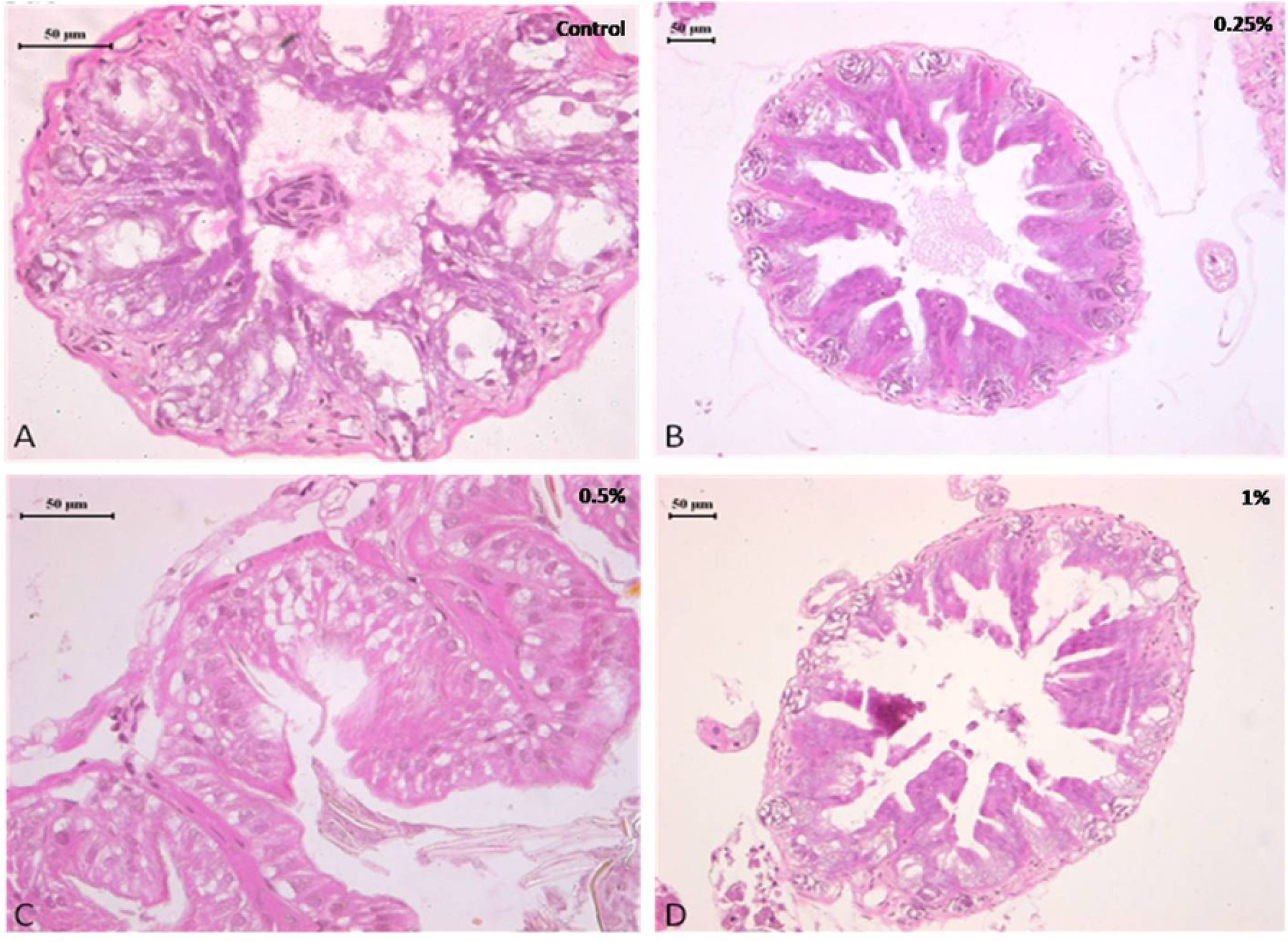
Photomicrograph of control and deltamethrin treated cockroach mid gut: A, Control; B, treated with 0.25% deltamethrin; C, treated with 0.5% deltamethrin; D, treated with 1% deltamethrin.

#### Deltamethrin quantification

HPLC method could estimate the deltamethrin on the exposed body of *P. americana* during the experiments. It was found that the average deltamethrin was 0.21±0.05 mg/cockroach (95% CI - 0.18-0.25; CoV 23.9%) (Supplementary File 2 showing median weight of exposed cockroaches and deltamethrin extracted using HPLC). The HPLC chromatograms of insecticide mixture (including deltamethrin) used as standard and deltamethrin extracted from the *P. americana* surface has been shown in figure 5 (A&B). HPLC could not detect deltamethrin in the brain, gut and ovary tissue of the exposed cockroaches.

**Figure 5:**
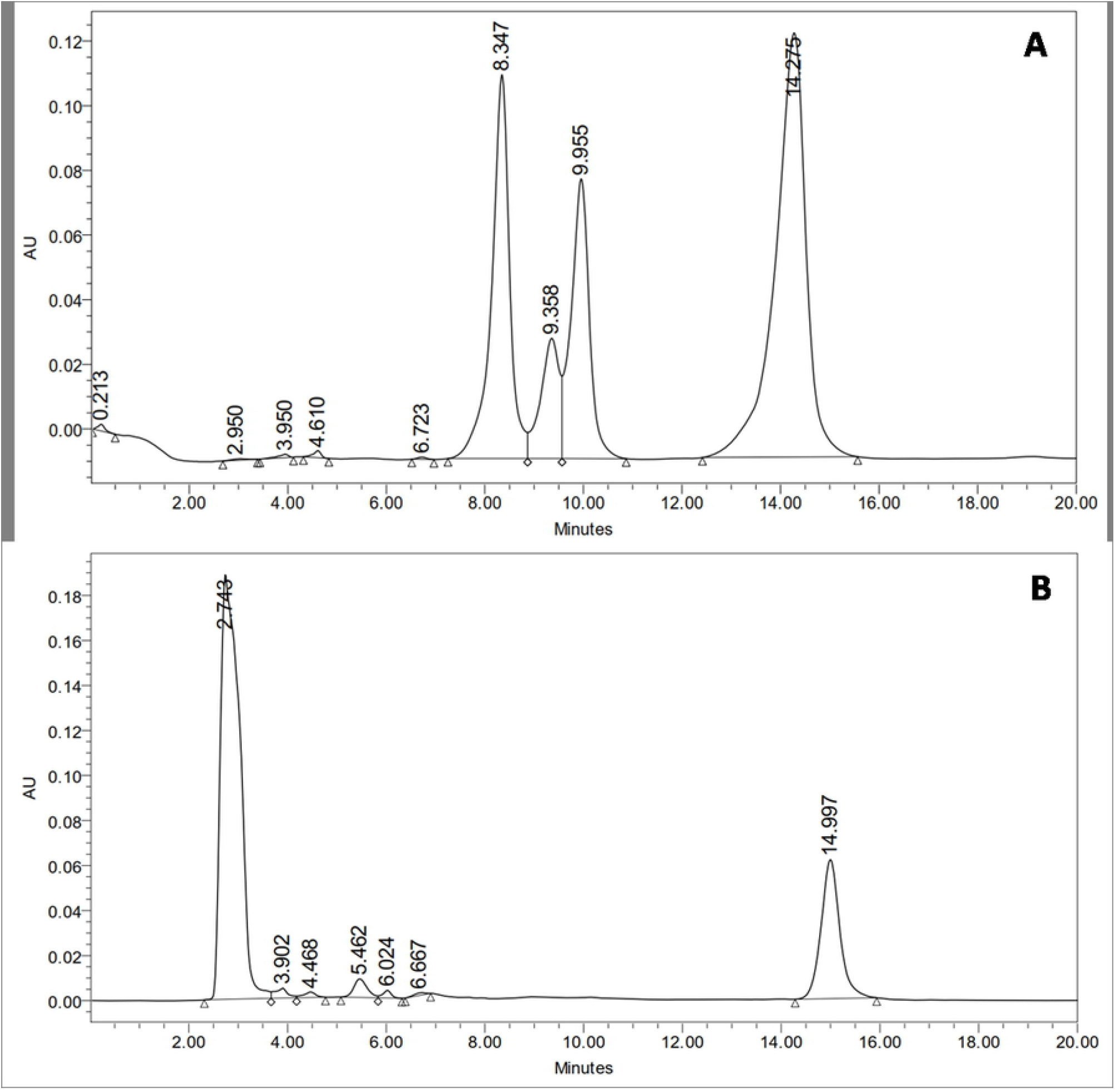
HPLC estimation of deltamethrin on *P. americana* exposed to 1% deltamethrin. (A) HPLC-UV chromatogram of insecticides mixture including deltamethrin used as standard; Pyriproxifen (9.358 min), Chloropyriphos (9.995 min) and Deltamethrin (14.275 min); (B) HPLC-UV chromatogram for deltamethrin extracted from whole cockroach; Deltamethrin (14.997 min).

## Discussion

Both *P. americana* as well as *B. germanica* are extremely common and among the most adapted detrimental pests that survive almost any environment around human [3,22](Bonnefoy et al. 2008; Fakoorziba et al. 2010). Besides being vectors of commonly occurring human pathogenic bacteria, cockroaches have been found of hosting many antibiotic-resistant bacteria in their digestive system [23](Fardisi et al. 2019). Control of cockroaches may be difficult due to several reasons; including development of resistance to repeatedly used insecticides, nevertheless resistance impact can be minimized through pre-assessment of efficacy in addition to rotational use of different insecticides.

The present research has been focused on evaluating deltamethrin mediated toxicity (including knock-down, delayed mortality and tissue level degeneration in critical internal organs) through time bound contact exposure to understand the effectiveness of deltamethrin as active ingredient (AI) in controlling cockroaches. Both the species showed high knock-down sensitivity to the insecticide as the knock-down ability was reduced ranging from 66.7% (at 0.00025%) to 100% (at 1%) during the short term continuous exposure of 1 hr. The results have suggested the concentration dependent knock-down and 24 hr delayed mortality in both the tested cockroach species. Therefore the LD values were found comparatively higher when exposure was given for lower time period. Both the species displayed about 60% survivility for 24 hr holding while about 30% survivility for 72 hr holding at lowest concentration. On the other hand 100% reduction in tested species was recorded at the concentration above 0.25%. This shows that probably sufficient amount of insecticide could not be absorbed through the cuticle at lower concentration.

Variety of insecticides has been evaluated against cockroaches and many of them have been found effective at low concentrations. A study conducted in Iran has shown that cockroach strains not much exposed to synthetic pyrethroids displayed >80% mortality as compared to those which were exposed due to irregular use of insecticides, mainly pyrethroids [24](Shahi et al. 2008). Synthetic pyrethroids have been shown to reduce the population of German cockroaches by >80% in India by the first week of treatment [25](Agrawal et al. 2005). Although the toxic effects are inherent capability of an insecticide and largely vary among different species of insects and even similar but geographically isolated species due to various reasons [26,27](Dhiman et al. 2013; Yadav et al. 2015), however studies have reported topical application remains the most sensitive method to ascertain the effectiveness [28](Holbrook et al. 2003). Many studies have demonstrated moderate to high level of resistance in cockroach species to different insecticides. Chai and Lee [29](2010) have reported heavy resistance for deltamethrin and cypermethrin against both *P. americana* and *B. germanica* in Singapore. In another study it was found that there was only 20% reduction in number when cockroaches were exposed to different pyrethroids [30](Fardisi et al. 2017). The resistance to insecticides in cockroaches develops faster probably because these inhabit relatively closely in large populations unlike mosquitoes and bugs, hence experience high level of selection pressure. In the present study, both the tested species were sensitive to deltamethrin and recorded >96% mortality in 0.025% deltamethrin suggesting that the deltamethrin resistance levels are extremely low in the test populations. Fardisi et al. [23](2019) has reported that against cockroaches an insecticide can be effective to reduce the populations in an areas having low starting resistance due to limited or no historical exposure of insecticides. Furthermore the study also suggested that the insecticides may exert repellency effect if fail to produce mortality due to increased resistance level.

Present results suggested that short term (24 hr) holding post 1 hr exposure of 0.25% and 0.5% deltamethrin did not cause considerable degeneration in the brain and ovary tissue. The brain showed normal morphology with distinct neuron cell bodies. It has been well documented that synthetic pyrethroids normally damage the nerve cells [31,32](Gutierrez et al. 2016; Soderlund, 2012), but no such damage was found in the present study. Furthermore the ovary also displayed normal panostic architecture with distinct oocytes and ovarian follicles without any fissure. Insecticide indoxacarb has been found to cause malformation in the oocytes displaying abnormal yolk and vacuolated follicular epithelium [33](Razik and Abd El-Raheem, 2019). However, similar to the present results, pyrethroid lambda-cyhalothrin exposure did not show any considerable alteration in the ovarian structure [33](Razik and Abd El-Raheem, 2019). Nevertheless exposure to 1% deltamethrin caused degeneration in gut epithelium affecting overall cytomorphology. This suggested that the survival deficits observed might have been resulted due to considerable disturbance in the physiological processes such as cellular metabolism and signal transmission in addition to the tissue damage. The sections did not show cytoplasmic granulation indicating that short time exposure was not enough to allow accumulation of deltamethrin or its by-products in the tissue system. However it has been suggested that 24 hr continuous exposure of deltamethrin caused its bio-accumulation in the midgut cytoplasm of *C. radiatus* nymph in the form of basophilic granules [31](Gutierrez et al. 2016). The vacuolization in the midgut could be initiation of deltamethrin mediated degeneration process. In an earlier study it was observed that cytoplasmic vacuolization was associated with deltamethrin-mediated initial changes were, which resulted in autophagy and apoptosis [31](Gutierrez et al. 2016). The toxic effect of deltamethrin may not necessarily be due to direct toxicity but deltamethrin exposure may result homeostasis imbalance in the midgut thereby reducing the ability to digest food for nutrients [34](Huang et al. 2015), dysregulation of the breathing activities [35](Unkiewicz-Winiarczyk et al. 2012) and also limiting in the production of signaling molecules that regulate its own physiology [36](Caccia et al. 2019). Deltamethrin was detected on the body surface of cockroach but not found in brain, ovary and gut tissue using HPLC. This suggests that deltamethrin up-taken through oral or cuticular route in the used concentrations was not sufficient enough to reach and accumulate in the internal tissues.

Resistance to insecticides continuous to exacerbate the impact of cockroaches on public health and overall hygiene. It is obvious that resistance to one or another class of insecticides is ubiquitous among the cockroach populations due to use of different insecticides based products. Still it may not be logical to simply guess the level of resistance at any place. Therefore efficacy assessment of an insecticide, because of its ability to identify suitable insecticide, is vital for choosing an appropriate insecticide for control success. Deltamethrin, at present, is among the most used insecticides in different intervention programmes and was able to provide the considerable efficacy against two cockroach species in the present study.

## Conclusion

Low concentration of deltamethrin was effective against two important field collected laboratory reared cockroach species. Even short term exposure displayed immediate knock down, delayed mortality and considerable histological damage in gut of tested species. The resistance to deltamethrin may be widespread among various public health importance insects due its use in various “over the counter” available products, however the suitable formulations such as insecticide paints, attractant baits etc. made using deltamethrin as active ingredient could be useful in the control operations.

## Acknowledgements

Authors are thankful to Director, Defence Research Development Establishment, Gwalior, for supporting the work. The help of technical staff during the experiments is acknowledged.

## Authors’ contributions

SD: Developed the study idea, performed laboratory experiments, and prepared the manuscript. KY: Involved in dissection, culture maintenance, performed the experiments. BN: Involved in HPLC estimation and drafted the manuscript. RRG: HPLC experiments. DPN: Histological assessment and manuscript editing. All authors read and approved the manuscript.

## Funding

No specific fund was received for this study.

## Availability of data and materials

All relevant data on the study are available from the corresponding author on reasonable request.

## Declarations

### Ethics approval and consent to participate

Not applicable.

### Consent for publication

All authors have given consent to publish this manuscript.

### Competing interests

The authors declare no competing interests

## Figure Legends

**Supplementary File 1:** Whisker Plot figure for weight (g) of adult *P. americana* exposed to 1% deltamethrin and deltamethrin uptake (mg) as estimated using HPLC.

## Notes

### Competing Interest Statement

The authors have declared no competing interest.

